# Using Virtual Patients to Predict Perceptual Outcomes for Optogenetic Sight Recovery Technologies

**DOI:** 10.64898/2026.04.09.716230

**Authors:** Vaishnavi B Mohan, Ezgi Irmak Yucel, Ione Fine, Geoffrey Boynton

**Affiliations:** Department of Psychology, University of Washington, Seattle, WA

**Keywords:** Optogenetics, Perception, Vision Restoration, Psychophysics, Retinitis Pigmentosa, Macular Degeneration, Visual prostheses

## Abstract

Optogenetics is emerging as a powerful approach for partial vision restoration, with at least three ongoing clinical trials in humans testing novel light-sensitive proteins (opsins) in patients with inherited retinal degenerative disorders. These therapies aim to restore light responsiveness by introducing opsins into surviving retinal cells, such as bipolar or ganglion cells, enabling them to generate neural activity in response to visual stimuli. One ongoing difficulty in selecting promising opsins for clinical development is that there is no way to predict patient perceptual outcomes from optogenetically evoked neural activity as measured *ex vivo*. Here, we introduce a *virtual patient* framework that quantitatively links the sensitivity and speed of opsin-mediated retinal responses to predicted patient outcomes, and show how this framework can predict temporal contrast sensitivity functions - a well-established measure of perceptual performance - from microbial opsin photocurrent responses. Our simulations demonstrate that opsin sensitivity and kinetics jointly determine perceptual outcomes, and that enhancing sensitivity at the expense of temporal resolution can degrade the perception of fast-moving stimuli. This computational platform provides a generalizable tool for comparing and selecting the most effective opsins for clinical translation, thereby guiding the design and optimization of next-generation sight restoration strategies.

## Main

Currently, over 200M people worldwide^1^ suffer from retinal neuro-degenerative disorders, such as retinitis pigmentosa and macular degeneration, diseases in which the photoreceptors in the retina progressively degenerate, leading to partial and/or complete loss of vision at later stages of the disease. Optogenetic sight restoration technologies aim to restore some vision in patients with these diseases by introducing the gene for a light-sensitive protein (opsin) to surviving bipolar or ganglion cells via carrier viral vectors. Upon photoactivation, the expressed protein undergoes a conformational change leading to neuronal responses (either depolarization or hyperpolarization) within target retinal cells, thereby making them light-sensitive and restoring partial vision^2,3^. Two recent clinical trials have reported partial sight recovery in humans^4,5^, with several more trials underway^6–10^, exploring a variety of opsins.

At present, selecting promising opsins for clinical trials is complicated by the lack of any quantitative way to predict perceptual effectiveness in patients from *ex vivo* photocurrent responses. As a result, optogenetic sight recovery technology relies on ‘best-guessing’ the perceptual efficacy of a given opsin, and human trials are initiated with substantial uncertainty as regards best-case efficacy as well as safety. This level of uncertainty is both ethically and practically problematic, given that opsin delivery is largely irreversible and human clinical trials often cost tens of millions of dollars.

Here we describe a computational ‘virtual patient’ framework that links opsin-mediated neural responses to ‘best-possible’ predicted perceptual outcomes, thereby enabling prediction of the ‘best possible clinical perceptual outcome’ of any opsin based on its response to photoactivation *ex vivo*. Using this framework, we show how the temporal dynamics of three representative microbial opsins (ChR2, ReaChR, and ChRmine) result in distinct perceptual outcomes. This predictive preclinical framework for estimating perceptual outcomes provides a powerful method for informed selection of promising opsins prior to clinical translation.

It is known that the effectiveness of opsins in sight recovery is likely to be heavily determined by two key biophysical properties: sensitivity and response kinetics^11^. As far as sensitivity is concerned, the maximum and sustained photocurrents generated upon photoactivation vary widely across opsins^12^. Sensitivity also varies as a function of wavelength: Opsins differ both in the wavelength of light corresponding to peak sensitivity, and the breadth of their wavelength tuning^13,14^. Finally, sensitivity is also likely to be monotonically (but not linearly) related to expression levels^15–17^.

Opsins also differ dramatically in their response kinetics - how quickly they open (*τ*_*on*_), reach steady state (*τ*_*steady*_), and close (*τ*_*off*_) the photo-activable pore upon light activation and cessation. These response kinetics determine the ability of opsin responses to faithfully follow luminance fluctuations at higher temporal frequencies, which in turn, determines the temporal sensitivity of retinal responses^18^.

## Results

All code was implemented in MATLAB and is publicly available on our [GitHub repository](https://github.com/Vaishnavi-B-Mohan/p2p_optoExpt.git). The framework is fully modular and supports modeling perceptual outcomes across a broad range of microbial opsins and visual stimuli.

### The relationship between opsin-mediated photocurrent responses and temporal sensitivity

The responses of opsins to light onsets and offsets are well described by the four-state photocurrent model proposed by^19^. Fig.1A shows simulated neural photocurrent responses for three microbial opsins with distinct sensitivity and kinetic profiles^20^. Microbial opsins exhibit a characteristic large transient depolarization current at stimulus onset (*τ*_*on*_, time taken to reach 63.2% of the maximum response). This peak response declines to a lower steady-state current during sustained illumination (*τ*_*steady*_, time taken to reach steady state), and declines again to the deactivated state upon stimulus offset (*τ*_*off*_), see Table 1. Although all three time constants play a role, for most opsins, once light adapted to a steady state, *τ*_*off*_ is the longest time constant, and is therefore the main rate limiting factor that influences temporal resolution, as shown in Fig. 1B. Opsins with rapid kinetics have less time to integrate photons, and therefore tend to be less sensitive, as shown in Panel C.

**Table 1:**
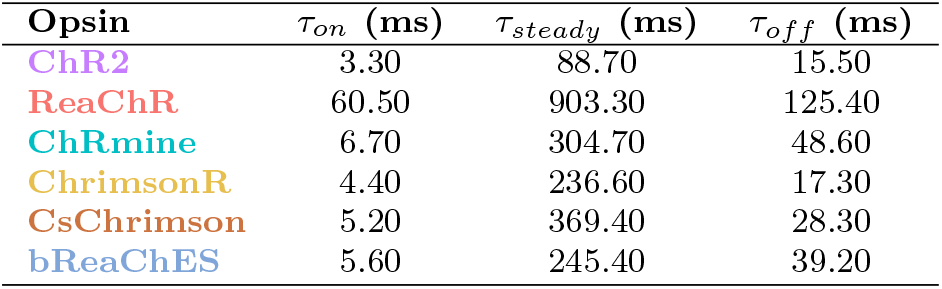
Opsin Time Constants. *τ*_*on*_: time taken for the opsin to open ion channels upon stimulus onset. *τ*_*steady*_ : time taken for the opsin photocurrent to reach a steady state from its initial transitionary peak photocurrent for a constant stimulus. *τ*_*off*_ : time taken for the photocurrent to decay after stimulus offset.

| Opsin | $\tau_{on}$ (ms) | $\tau_{steady}$ (ms) | $\tau_{off}$ (ms) |
| --- | --- | --- | --- |
| ChR2 | 3.30 | 88.70 | 15.50 |
| ReaChR | 60.50 | 903.30 | 125.40 |
| ChRmine | 6.70 | 304.70 | 48.60 |
| ChrimsonR | 4.40 | 236.60 | 17.30 |
| CsChrimson | 5.20 | 369.40 | 28.30 |
| bReaChES | 5.60 | 245.40 | 39.20 |

**Table 2.** Parameters of the four-state photocurrent opsin model ^20^.

| Parameter | ChR2 | ReaChR | ChRmine |
| --- | --- | --- | --- |
| $G_{d1}$ ($\text{ms}^{-1}$) | 0.09 | 7.7e-3 | 0.02 |
| $G_{d2}$ ($\text{ms}^{-1}$) | 0.01 | 1.25e-3 | 0.013 |
| $G_r$ ($\text{ms}^{-1}$) | 5e-4 | 3.33e-5 | 5.9e-4 |
| $g_0$ ($\text{mS cm}^{-2}$) | 0.12 | 0.28 | 2.2 |
| $\Phi_m$ (photons $\text{mm}^{-2} \text{s}^{-1}$) | 4e16 | 5e17 | 2.1e15 |
| $k_{a1}$ ($\text{ms}^{-1}$) | 3 | 1.2 | 0.2 |
| $k_{a2}$ ($\text{ms}^{-1}$) | 0.18 | 0.01 | 0.01 |
| $k_f$ ($\text{ms}^{-1}$) | 0.03 | 0.012 | 0.001 |
| $k_b$ ($\text{ms}^{-1}$) | 0.003 | 0.001 | 0 |
| $G_{0,a1}$ ($\text{ms}^{-1}$) | 0 | 0 | 0 |
| $G_{0,a2}$ ($\text{ms}^{-1}$) | 0 | 0 | 0 |
| $G_{0,f}$ ($\text{ms}^{-1}$) | 0.015 | 5e-4 | 2.7e-4 |
| $G_{0,b}$ ($\text{ms}^{-1}$) | 0.005 | 5e-4 | 5e-4 |
| $\gamma$ | 0.05 | 0.05 | 0.05 |
| $p$ | 1 | 1 | 0.8 |
| $q$ | 1 | 1 | 1 |
| $E$ (mV) | 0 | 7 | 5.64 |
| $\lambda$ (nm) | 460 | 590 | 590 |

**Fig. 1:**
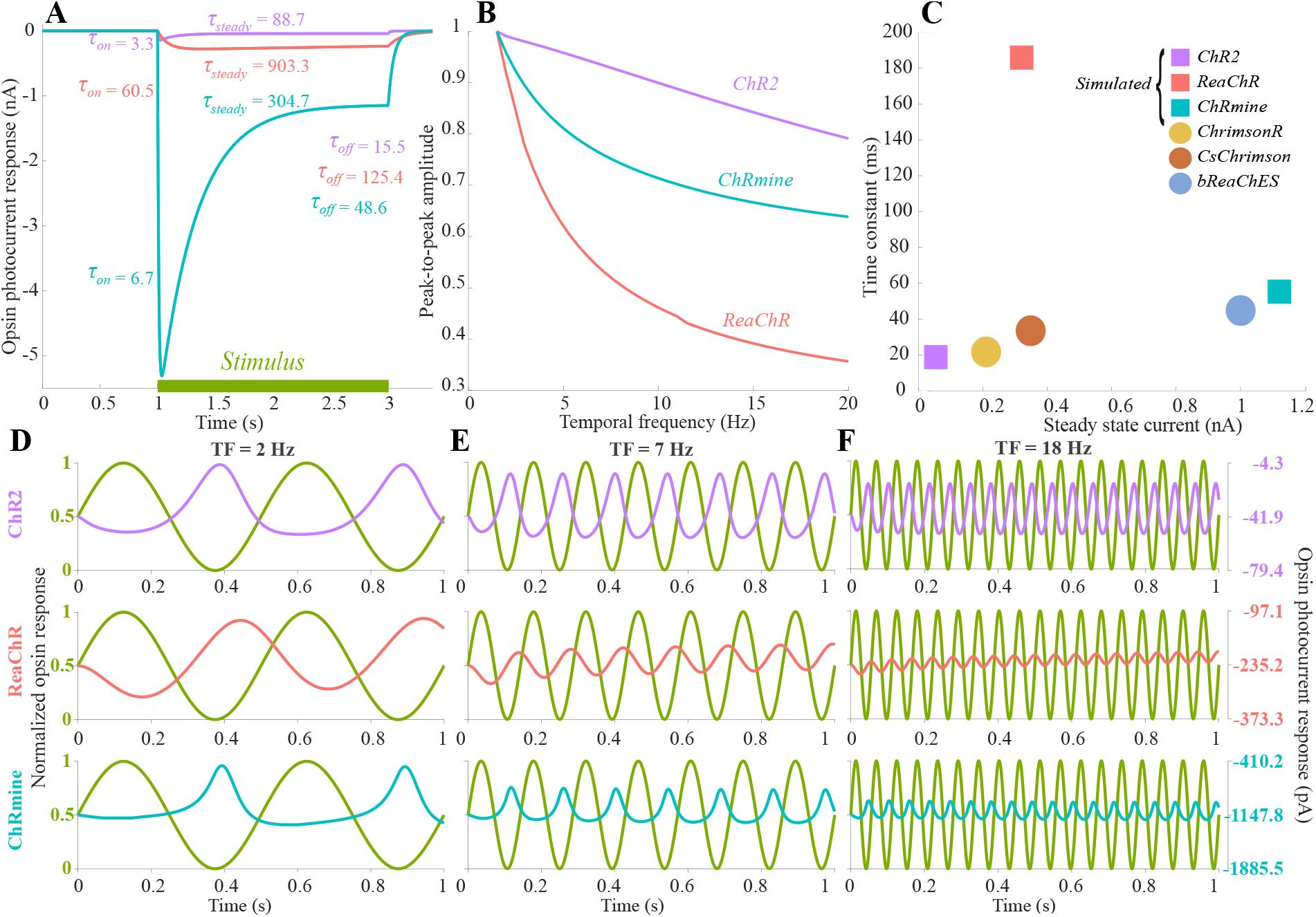
Comparison of opsin sensitivity and kinetics. Opsin photocurrents, simulated using the four-state photocurrent model^19,20^, highlight the inherent trade-off between opsin sensitivity and response kinetics for the three modeled opsins: ChR2, ReaChR, and ChRmine. **(A)** Opsin photocurrent responses for a constant stimulus (1 mW mm^−2^, 2 s) at each opsin’s peak wavelength (460 nm for ChR2; 590 nm for the others) highlight how sensitivity varies across the three opsins. **(B)** Normalized photocurrent responses are modeled for sinusoidal stimuli of amplitude (1 mW mm^−2^, 1 s) and varying temporal frequencies, after light adaptation. Y-axis indicates peak-to-peak amplitude of opsin photocurrent response and x-axis indicates the temporal frequency of the sinusoidal stimulus, highlighting how peak-to-peak amplitude drops with increasing temporal frequency for these three opsins. **(C)** Trade-off between opsin sensitivity and opsin dynamics. The x-axis represents the steady state photocurrent (in nA) to constant illumination (1 mW mm^−2^, 2 s) at the peak wavelength for each individual opsin and y-axis represents *τ*_*off*_. The model highlights the trade-off between steady-state photocurrent amplitude (nA) and offset response time constant (ms). **(D–F)** Simulated photocurrent responses for the three opsins to sinusoidal stimuli (normalized to a maximum amplitude of 1) at temporal frequencies of 2 Hz, 7 Hz, and 18 Hz, respectively. y-axis labels on the left and right indicate normalized contrast levels and absolute photocurrent values (pA) respectively.

For any stimulus timecourse it is possible to simulate opsin-mediated photocurrents *I*(*t*) by numerically solving the four-state photocurrent model (see Eqns. 1-4, in Methods) using Euler’s method. The lower panels of Fig.1 show predicted responses for our three modeled opsins (based on the four-state photocurrent model^19^) to sinusoidal inputs as a function of temporal frequency. Note the differences in y-axis scaling across the three opsins. While all these opsin responses preserve the periodicity of the input signal, the output waveforms are non-sinusoidal, reflecting a temporally nonlinear transformation (Fig.1D-F).

#### (i) ChR2

Channelrhodopsin-2 derived from the green algae *Chlamydomonas reinhardtii*, was used in the seminal study^3^ demonstrating optogenetic restoration of light sensitivity in RGCs. ChR2 has also been incorporated in a recently concluded human clinical trial NCT02556736. Among the three opsins considered here, ChR2 exhibits the fastest kinetics but the lowest sensitivity. In addition, its sensitivity peaks in the blue region of the visible spectrum (460 nm)^21^. This short-wavelength peak, combined with low sensitivity, makes it difficult to elicit robust photocurrents within retinal light intensity safety limits^22^, even when using light intensifying goggles. Most engineered microbial opsins used in vision restoration are derived from ChR2, incorporating modifications to make them more sensitive and/or shift peak sensitivity to higher wavelengths. Losses in sensitivity as a function of temporal frequency can be seen by comparing Fig.1 Panels D-F (in **purple**). Because ChR2 has fast dynamics, there is still a robust response at 18 Hz. The response is also highly asymmetrical, with much stronger responses to luminance decrements than to increments.

#### (ii) ReaChR

ReaChR is a red-shifted variant of ChR2 with peak sensitivity near 590 nm and improved light sensitivity^23^. Under bright daylight conditions ReaChR is ∼ 6.25x more sensitive than ChR2. This increased sensitivity and longer peak wavelength makes it easier to elicit robust photocurrents while remaining within safe retinal illumination limits, but comes at the cost of substantially slower kinetics as compared to ChR2^24^. Because this opsin has slow dynamics, the response at 18Hz is heavily attenuated (Fig.1D-F, **red**). The response is less asymmetrical than that of ChR2, though responses to luminance decrements are still stronger than for luminance increments.

#### (ii) ChRmine

ChRmine is the most sensitive of the three opsins, ∼30x more sensitive than ChR2^25^. Like ReaChR, ChRmine exhibits peak sensitivity at longer wavelengths (∼590 nm), with kinetics between those of ChR2 and ReaChR. Since it has both high sensitivity and reasonably rapid kinetics, ChRmine is emerging as a promising candidate for clinical applications. While the first report of partial sight recovery in humans used ChRimsonR^4^, a less sensitive precursor, recent studies have demonstrated the viability of ChRmine in rd1 mouse retina models^26^. Because this opsin has intermediate temporal dynamics, the attenuation of the response at 18 Hz is intermediate between that of the other two opsins (Fig.1D-F, **blue**). The response is highly asymmetrical, with little response to luminance decrements.

### Predicting perceptual experience from opsin kinetics

Since the response of each opsin can be simulated for any visual stimulus, it is possible to predict patient percepts. We begin by simulating the experience of optogenetic vision using real-world videos (Fig. 2). Here, to focus on the impact of temporal dynamics, we scaled the opsins to have similar sensitivity at steady state. Despite scaling for sensitivity, perceptual outcomes are strikingly different across the three opsins. For the slower opsins, ChRmine and ReaChR, perceptual distortions are noticeable. Fast moving objects are heavily attenuated by motion streaks - e.g. in the bottom panel, the running pedestrian almost disappears for ReaChR. For the opsins with heavily asymmetric responses to increments vs. decrements (ChR2 and ChRmine) objects that are brighter than the surroundings (such as the buildings) are heavily attenuated; for example, in the top panel the white pants of the pedestrian blends into the light gray tarmac. Thus, it is possible to make qualitative predictions about patient perceptual experience by modeling the nonlinear temporal dynamics introduced by opsin kinetics. (One complication is that we view these movies through an intact visual system - compensation for this is discussed below.)

**Fig. 2:**
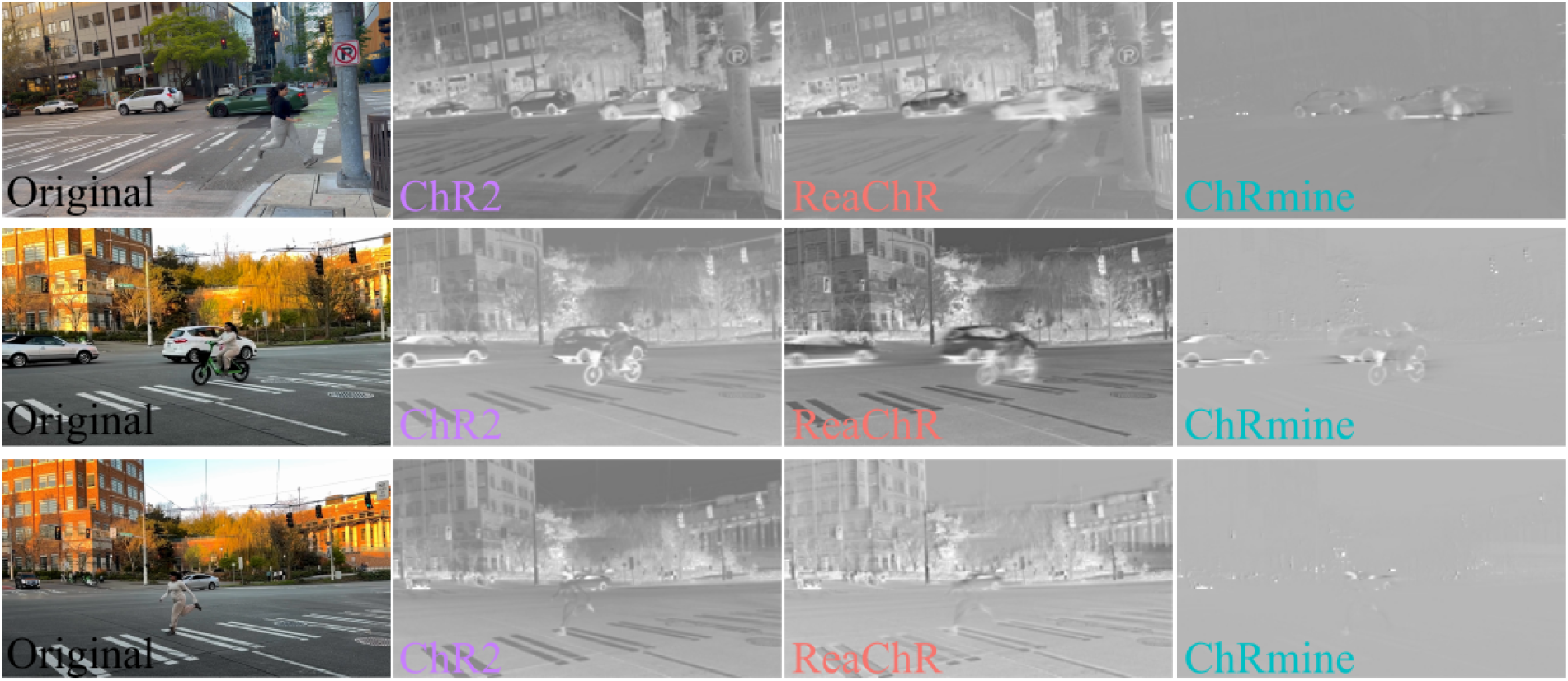
Simulated percepts from real-world movies demonstrate opsin-specific temporal distortions. Representative frames from real-world videos passed through the four-state photocycle model for three opsins. Responses were normalized to remove relative sensitivity differences. ChR2 closely mirrors neurotypical temporal dynamics and the resulting perceptual output does not differ greatly from a contrast-reversed image. ReaChR strongly attenuates higher temporal frequencies, suppressing fast-moving objects, including the pedestrian (top), the bike rider (middle), and the running pedestrian and silver car (bottom). ChRmine has intermediate kinetics, preserving the visibility of rapidly moving objects. However, ChRmine is highly non-linear, with very weak responses to contrast increments: as a result, objects that are brighter than the background are attenuated in contrast. *(Note: All video frames in this figure depict the authors of this study; no other individuals are shown. Full supplementary videos of the simulations are available from the corresponding author upon request.)*

### Human-in-the-loop Virtual Patients: Predicting clinical outcomes

Next, we extend this framework to quantitatively predict perceptual performance on standardized clinical tests (Fig. 3). Our example application is the temporal contrast sensitivity function (tCSF), but a similar approach could be used to predict performance on other standard clinical measures such as the Snellen Eye Chart.

**Fig. 3:**
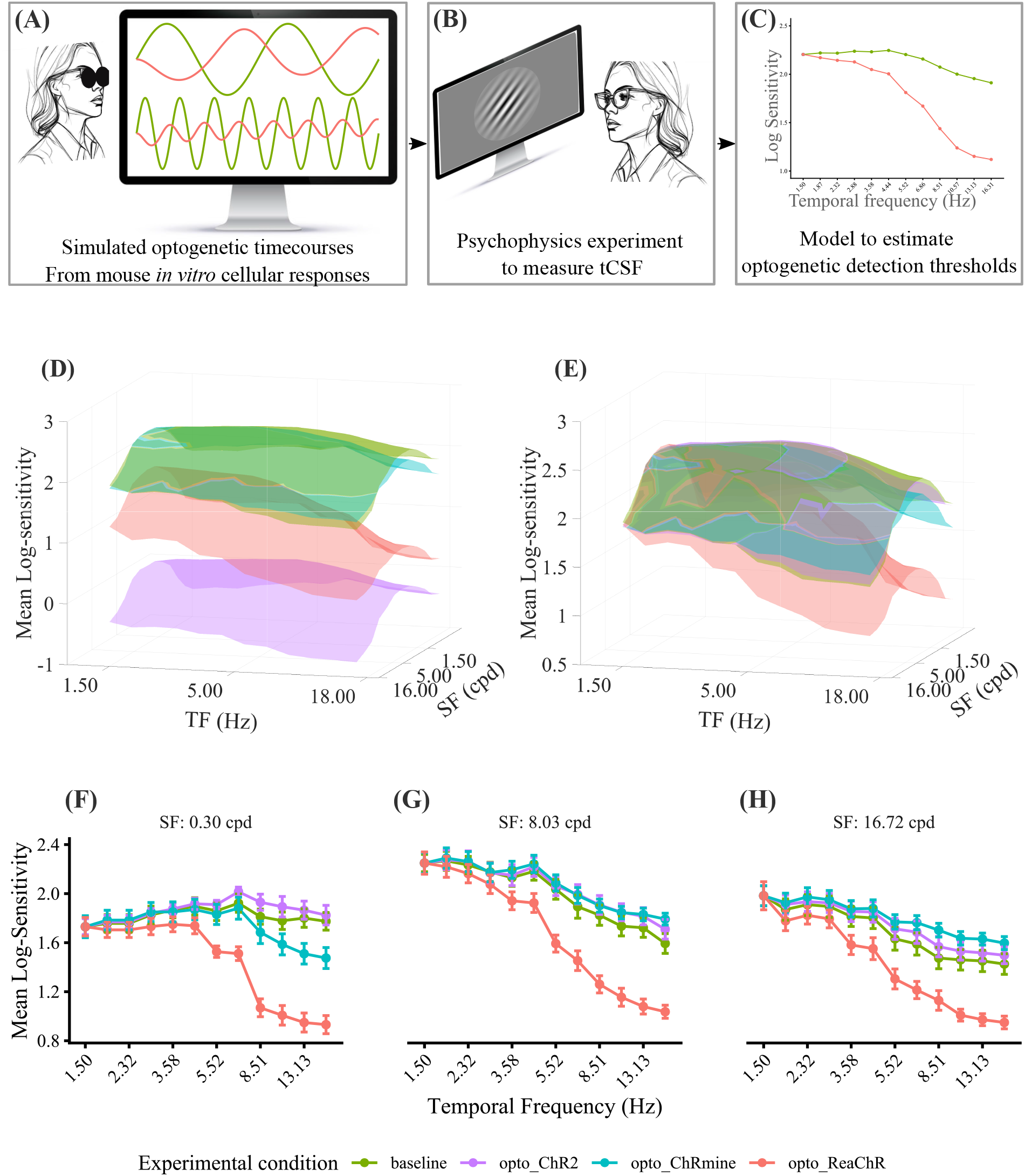
**(A-C) Flowchart of Virtual patient simulator.A**. Simulated optogenetics: A four-state photocycle model was used to describe the photocurrent response of each microbial opsin to any visual stimulus. **B**. Neurotypical participants (‘human-in-the-loop’) perform psychophysical judgments on stimuli designed to mimic the perceptual outcomes of optogenetic vision. **C**. This allows for estimates of best-case patient performance for clinical measures of visual performance. tCSFs for the unfiltered visual stimulus and the scaled tCSF for ReaChR opsin are shown. **(D-E) Simulated perceptual outcomes of optogenetic stimulation**. Temporal contrast sensitivity functions (tCSFs) derived from simulated responses in a two-alternative forced-choice (2AFC) orientation discrimination task are shown for neurotypical vision, ChR2, ReaChR, and ChRmine. These tCSFs are averaged across six virtual patients. **D**. Unscaled tCSFs reveal differences in perceptual sensitivity across opsins relative to control, where greater deviation from the neurotypical tCSF indicates larger losses in visual acuity. **E**. tCSFs normalized to the control’s peak sensitivity at the lowest temporal frequency highlight the influence of opsin kinetics. The slope of each curve reflects the rate of sensitivity decline with increasing temporal frequency, with steeper slopes indicating stronger attenuation of high-frequency visual signals. **(F-H) 2D tCSFs highlight perceptual effects of opsin kinetics**. 2D plots of normalized tCSFs (sliced from (E)) are obtained from psychophysics data across temporal frequency (Hz) observed at spatial frequencies: 0.30, 8.03 and, 16.72 cpd.

The time-courses of temporally modulated sinusoidal gratings were passed through our optogenetic model as described in Fig.1. These modified stimuli were then presented to sighted neurotypical participants in a ‘human-in-the-loop’ framework (illustrated in Fig. 3A). The psychophysical performance of these virtual patients - neurotypical participants viewing stimuli that replicate the experience of opsin mediated vision - represents the best-possible perceptual performance of patients transfected with that opsin.

tCSFs were estimated using a classic two-alternative forced-choice (2AFC) orientation discrimination task as shown in the virtual patient simulator schematic (Fig. 3B), which finds the lowest contrast needed to determine grating orientation (-45° or +45°) with 80% accuracy as a function of the spatial and temporal frequency of the stimulus (see Methods). Unmodified spatial sinuoidal gratings that were also modulated sinusoidally over time were used to establish neurotypical temporal contrast sensitivity functions (baseline tCSFs). These same stimuli, passed through our opsin models, were used to estimate opsin-specific tCSFs.

(Fig. 3D-E) shows tCSFs averaged across six individuals. Unscaled tCSFs (Fig. 3D) reveal substantial perceptual sensitivity differences across opsins, with a mean separation of 1.96 log units between ChRmine, the most sensitive opsin, and ChR2, the least sensitive. This corresponds to an approximately 94-fold difference in visual sensitivity.

To isolate the effects of opsin *kinetics*, we normalized the tCSFs for each opsin to match neurotypical sensitivity at the lowest temporal frequency (Fig. 3E). 2D plots showing 0.30, 8.03, and 16.72 cpd are shown in Fig. 3F-H. After normalization, ChR2 closely overlaps with the neurotypical tCSF, suggesting that under conditions of very high illumination ChR2 could support percepts with minimal temporal distortion. In contrast, ReaChR, the slowest opsin, strongly suppresses temporal frequencies above 8 Hz, with a loss in sensitivity of 1.5 log units at 16 Hz. This corresponds to an approximately 32 -fold reduction in sensitivity at higher temporal frequencies relative to the neurotypical tCSF. As shown in Fig. 2, this is likely to result in impaired perception of fast-moving stimuli. More sluggish opsins are also likely to be highly susceptible to motion streaks induced by eye-movements. Thus, the perceptual quality provided by an opsin depends on both sensitivity and temporal dynamics.

One important difference between virtual and real patients is that our virtual patients view stimuli through their intact outer retina. It is possible to compensate for this using an inverse temporal filter^27^ (see Methods). Compared to the dynamics of the opsins examined here, the effect of compensating for photoreceptor dynamics was negligible, as shown in Figure 4, but this step may prove more important as opsins improve.

**Fig. 4:**
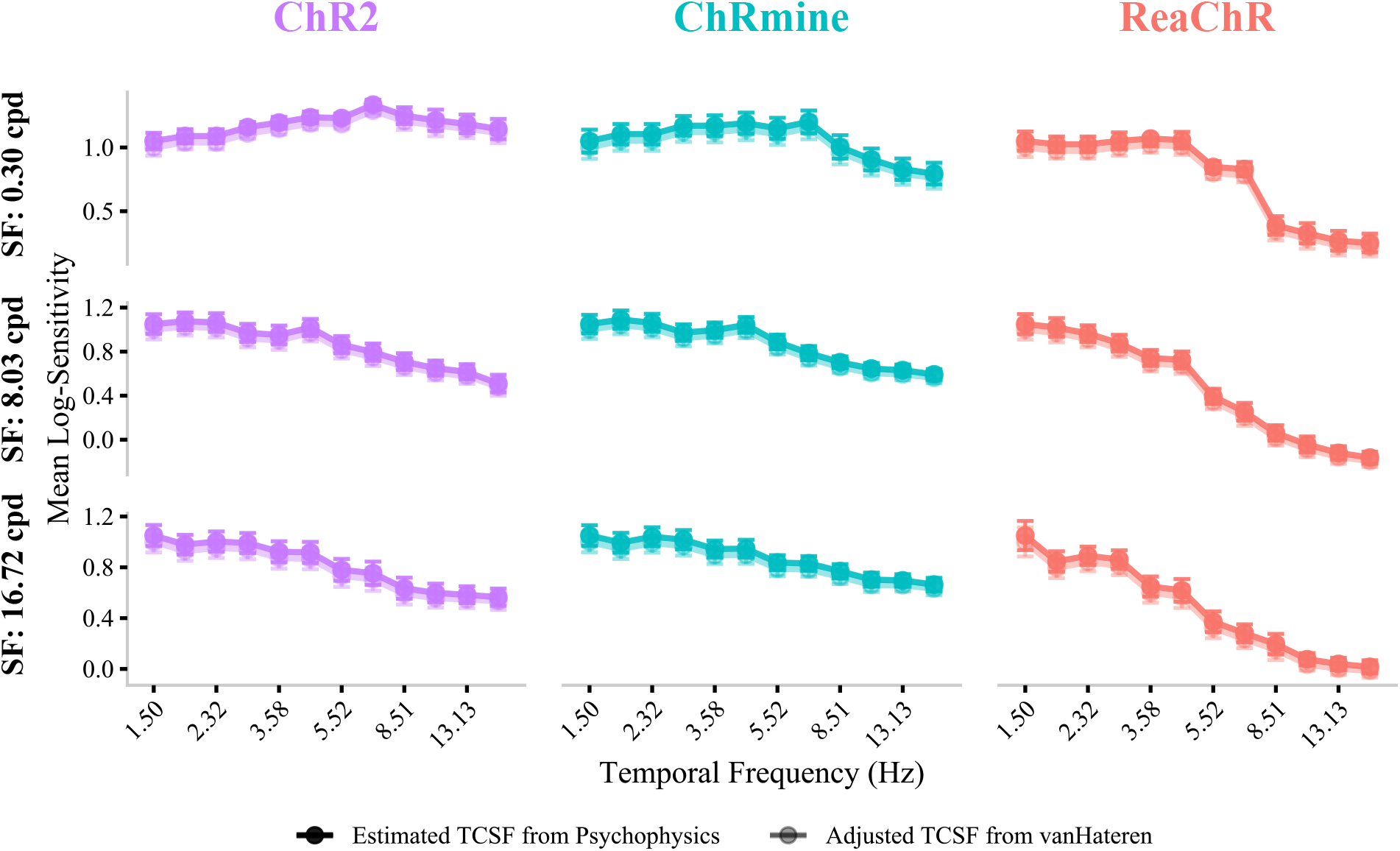
Compensating for photoreceptor dynamics in virtual patients. Comparison of contrast sensitivity with temporal frequency with human-in-the-loop Virtual Patients (solid colors) and adjusted sensitivities after correcting for photoreceptor processing (translucent colors), at three spatial frequencies for the three opsins. Human-in-the-loop contrast sensitivity functions (tCSFs) were generated by averaging psychophysics tCSFs obtained across six participants viewing gratings filtered by opsin time-courses. Correction was carried out by dividing the original tCSF values by the frequency response of photoreceptors based on the van Hateren model (shown in Extended Data, Fig.7) to obtain a better estimate of tCSFs corresponding to real optogenetic patients.

### Simulated Virtual Patients: Predicting clinical outcomes directly from opsin *ex vivo* kinetics

While our ‘human-in-the-loop’ psychophysical experiments with virtual patients can estimate best-possible performance across a wide variety of clinical tests, they are time-consuming, and therefore unsuitable for rapidly evaluating and comparing a large number of candidate opsins. To address this constraint, we developed a linking hypothesis model capable of predicting the temporal contrast sensitivity function (tCSF) of any microbial opsin directly from a dynamic model of its neural photocurrent response. We assumed (see Methods) that detection sensitivity is linearly related to rectified summed opsin responses over time (inspired by^28^), that detection sensitivity differs for positive (*β*_*inc*_) and negative (*β*_*dec*_) excursions from the mean response, and that detection sensitivity varies with the spatial frequency of the stimulus. Across all three opsins, we estimated *β*_*inc*_ and *β*_*dec*_, for each spatial frequency (see Extended data, Table 4) by minimizing the mean squared error between the psychophysical measurements corresponding to the three microbial opsins and model predictions of sensitivity. This provides an estimate of losses in detection sensitivity as a function of temporal and spatial frequency due to opsin’s kinetics, relative to the neurotypical CSF. As shown in Fig. 5 for three example spatial frequencies, fully simulated virtual patient tCSFs closely predict the performance of our human-in-the-loop virtual patients. Importantly, the same values of *β*_*inc*_ and *β*_*dec*_ fit all three opsins, allowing this model to be generalized to novel opsins without the need for additional data.

**Fig. 5:**
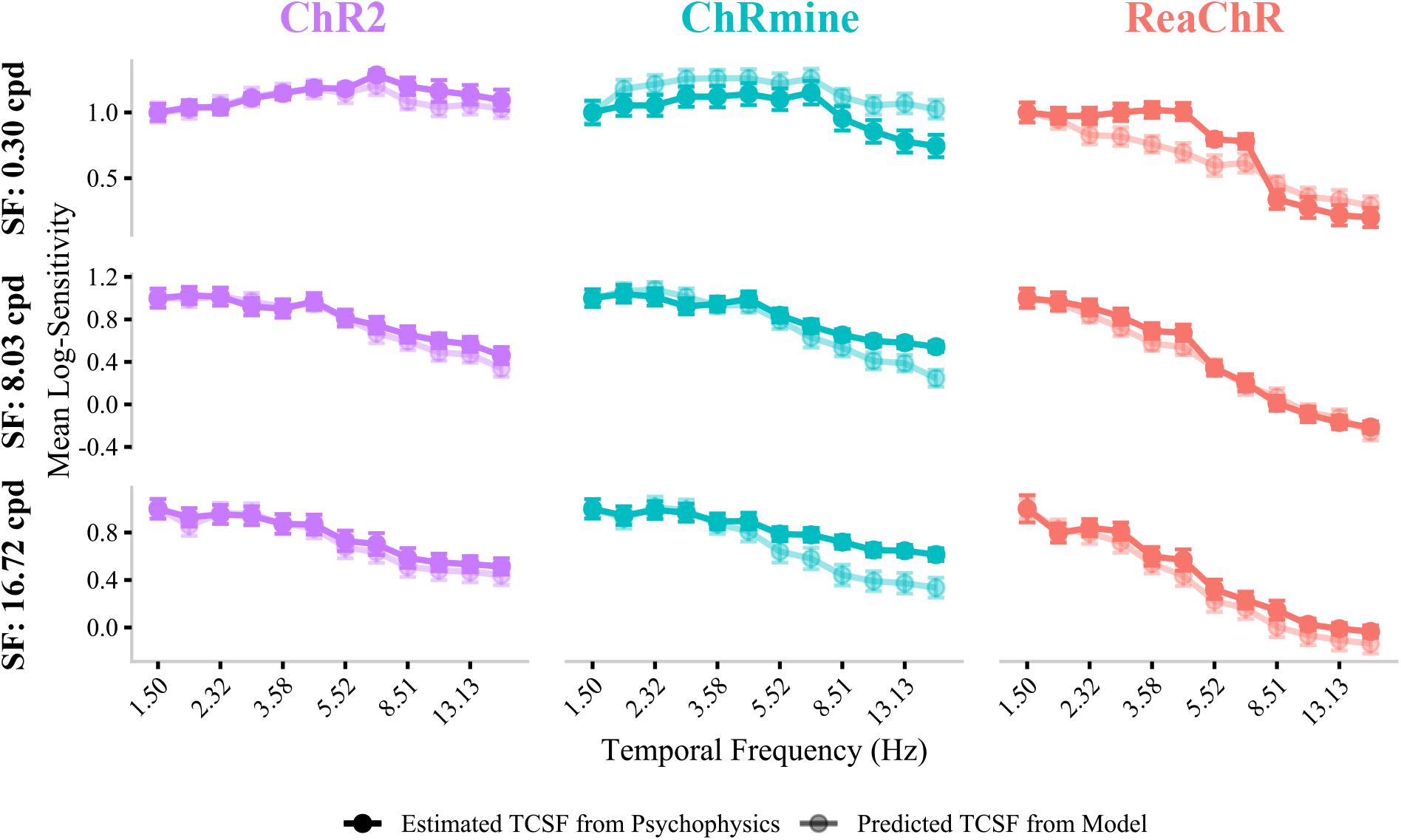
Predicting opsin tCSFs directly from *ex vivo* kinetics. Comparison of human-in-the-loop Virtual Patients (solid colors) and Simulated Virtual Patients (translucent colors). Human-in-the-loop temporal contrast sensitivity functions (tCSFs) were generated by averaging across six participants viewing gratings filtered by opsin time-courses (replotted from Fig.4, solid colors). Simulated Virtual Patients tCSFs were generated from the neurotypical tCSF for each individual. Threshold contrast gratings were filtered by opsin time-courses, and the linking hypothesis model was then used to predict probability correct as a function of temporal and spatial frequency. Error bars represent the standard deviation across subjects.

Thus, adding a simple linking model that converts visual input into behavioral performance creates a rapid and generalizable approach for approximating clinical perceptual outcomes for any microbial opsin based solely on its neural photocurrent response to visual stimuli.

## Discussion

Opsin selection for human optogenetic sight recovery currently relies on guessing which photocurrent profile will be most efficacious. Certainly, an opsin must be adequately sensitive. For example, ChR2, which has temporal dynamics that closely mirror neurotypical temporal responses (Fig. 2), has low sensitivity that does not seem to be capable of eliciting useful neural visual responses under natural illumination. Clinical trial NCT02556736 reported no difference in full-field sensitivity thresholds between treated and untreated eyes^29,30^. As a result, sensitivity is currently prioritized when selecting opsins. However, the functional benefits of increasing sensitivity may diminish once a threshold level of sensitivity has been achieved, and this threshold may be well below the sensitivity of natural vision. The visual system operates without any major loss of function over approximately 10 log levels of retinal illumination, and up to 40-50% of ganglion cells in the periphery or about 25-35% in the central retina can be lost before visual sensitivity losses exceed the 95% confidence limits^31,32^. Thus, one might expect some degree of robustness to both reduced levels of activation and a subset of cells being transfected.

In contrast, our virtual patients reveal that even relatively subtle differences in temporal kinetics and asymmetric response profiles can play an important role in shaping visual experience, suggesting that both these aspects of opsin selection may also play an important perceptual role^33^.

We describe here two types of virtual patients: the human-in-the-loop virtual patient and the fully simulated virtual patient. The ‘human-in-the-loop’ version relies on neurotypical observers viewing stimuli designed to replicate the experience of optogenetic vision. There are four important strengths to this ‘human-in-the-loop’ approach. The first is that it is possible to match virtual patients for important non-ophthalmological features that may influence clinical performance - such as age. The second is that our virtual patient provides a way to bridge the gap between well-controlled psychophysical measurements and assessing performance under conditions that genuinely reflect real-world outcomes. Currently, tests of visual restoration technologies tend to be ‘uneasy compromises’ where participants perform real world tasks under highly simplified visual conditions (such as differentiating between a small number of items on a plain background^4,34^, or navigating under highly simplified real world conditions^35^. Although these tests reflect real world tasks, they are carried out under conditions that do not reflect the cluttered dynamic conditions of natural vision. Our model of opsin dynamics is parallel in structure, and therefore could easily be converted into the virtual reality domain, making it possible to test the performance of human-in-the-loop virtual patients under conditions that genuinely mimic the demands of natural vision. A third, related, advantage of our approach is that it allows for more qualitative judgments - such as asking virtual patients which opsin provides the ‘best’ perceptual experience. Finally, it is possible to examine how well individuals can adapt to optogenetic vision with training, an approach that has previously been applied to electrical sight restoration technologies^36,37^.

The disadvantage of the ‘human-in-the-loop’ approach is that it is time-consuming and requires collecting human psychophyical data. We therefore extended our framework to include a linking hypothesis model to relate predicted visual experience to behavioral performance. This approach provides a highly efficient way to pre-screen microbial opsins to select promising candidates for more extensive behavioral testing.

Our simulations also suggest that the choice of opsin may depend on what functional benefits should be prioritized. An opsin with high sensitivity and slow kinetics might allow for reading in indoor conditions, but be less suitable for mobility tasks - fast-moving stimuli can become imperceptible with slower opsins. In contrast, an opsin with lower sensitivity but faster kinetics might offer little perceptual benefit indoors or at night, but might improve mobility under outdoor daylight conditions.

Because our model is based on the retinal image, incorporating the motion streaks induced by eye movements is a straightforward extension of our framework. We did not include eye movements in the simulations shown above because pilot simulations found that incorporating naturalistic eye movements had little impact on the predicted tCSFs. However, the interaction between opsin dynamics and eye movements is likely to play a more important role during natural viewing. Under typical conditions, saccadic suppression—lasting approximately 100-200 ms (depending on the amplitude of eye-movement) — prevents the perception of motion streaks on the retina^38^. Our simulations suggest that opsins with slower temporal dynamics (those showing significant attenuation at ∼3-5 Hz) could produce perceptually noticeable motion streaks, although there is evidence that saccadic suppression may be adaptable depending on visual context^39,40^.

We adjusted our tCSFs to compensate for the intact outer plexiform layer in our human-in-the-loop virtual patients based on a linear filter. When fully adapted, processing in the outer plexiform layer can be approximated as linear over a limited range of stimulus intensities. We exploited this approximation to construct an inverse temporal filter^27^ (see Methods), which we used to account for the temporal processing introduced by the photoreceptors. Compared to the dynamics of the opsins examined here, the effect of compensating for photoreceptor dynamics was negligible, and the linear model was therefore sufficient. However, accounting for non-linear photoreceptor dynamics may become more important as opsin performance improves^41–43^.

Although our framework provides a generalizable way to estimate preclinical tCSFs, it is important to note that the model does not predict actual patient performance. Rather, it reflects a “best-case” scenario—the upper limit of perceptual quality given the modeled factors. In practice, a range of additional factors are likely to result in perceptual outcomes that fall short of this idealized bound.

Our model does not account for cell-type–specific differences among retinal ganglion cells. Current optogenetic therapies indiscriminately confer photosensitivity to both ON and OFF RGCs, whereas cortical circuits rely on distinct functional signals from these pathways. Simultaneous activation of all RGCs appears to produce bright percepts, although the mechanisms underlying this effect remain unclear^41^. It is plausible that unselective stimulation reduces sensitivity. In the case of electrical stimulation, thresholds can decrease by up to an order of magnitude with extensive training^42,43^, suggesting that unselective activation is associated with a loss of sensitivity.

Finally, our current model assumes uniform opsin expression across the retina. In reality, spatial heterogeneity in expression density is likely to introduce perceptual distortions. Moreover, retinal layers undergo substantial reorganization following degeneration, which is likely to interact with opsin expression, creating additional spatial distortions. At present, no systematic model exists to predict these structural changes, but incorporating them in future work could provide important insights into how spatial patterns of opsin expression and neuronal reorganization shape perception.

In conclusion, we describe here an initial framework for predicting clinical outcomes for opsin technologies from *ex vivo* photocurrent data. Future directions should include elaborating this model to incorporate cell-type specificity, retinal degeneration, and opsin expression density. However, even in this early form, this work provides a useful method for systematically evaluating perceptual viability of optogenetic therapies before clinical application.

## Methods

In neurotypical vision, light entering the retina hyperpolarizes photoreceptors, leading to a discharge of stimulus-dependent photocurrents that modify the membrane potential of downstream retinal layers, triggering firing within retinal ganglion cells. In the case of opsin-mediated vision, visual stimuli evoke opsin-mediated photocurrents that modulate the firing rates of RGCs (or graded potentials in bipolar cells), resulting in a neural signal that is transmitted to the brain. As described above, the temporal response of opsin-driven photocurrents depends on the specific opsin and differs substantially from native human photocurrents observed in neurotypicals.

### 1 Four-state Photocurrent Model

In the case of microbial opsins, a well-established photocycle model proposes that the microbial opsin exists in four possible conformational states: two non-conducting closed (*C*_1_ and *C*_2_) and two conducting open (*O*_1_ and *O*_2_)^19^. It is hypothesized that light-evoked transitions can occur either under dark adaptation (*C*_1_ → *O*_1_) which is faster, or under the slower light-adapted (*C*_2_ → *O*_2_) condition. Transitions between open states are reversible and are caused by intrinsic conformational changes, while *C*_2_ → *C*_1_ is thermally induced.

*C*_1_, *C*_2_, *O*_1_ and *O*_2_ represent the fraction of opsin molecules in one of the four possible states at any given time *t*. Therefore, *C*_1_ + *C*_2_ + *O*_1_ + *O*_2_ = 1. The transition rate of molecules in each of these four states can be described by the following set of ordinary differential equations:

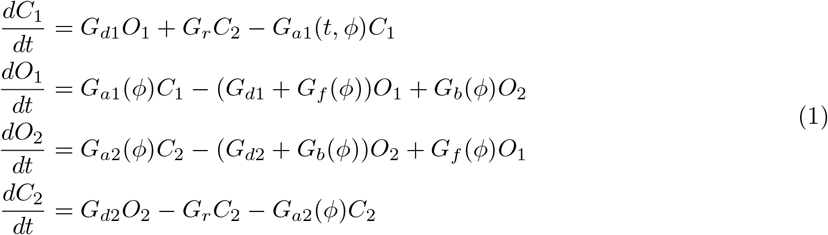

*G*_*a*1_ and *G*_*a*2_ denote the activation rates governing transitions from closed to open states, whereas *G*_*f*_ and *G*_*b*_ describe the forward and backward transitions between open states. Because transitions involving open states are driven by light, these rates depend on the photon flux per unit area (*ϕ*(*t*)) of the visual stimulus incident on the retina at time *t* given by:

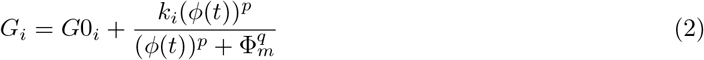

where, *i* ∈ [*a*1, *a*2, *f, b*]. Φ_*m*_ denotes the half-saturation photon flux, the value of photon flux at which a light-dependent transition reaches half of its maximum rate; lower values mean that the opsin is more sensitive to dim light. The stimulus-dependent photon flux per unit area incident on the retina is given by Eqn. 3:

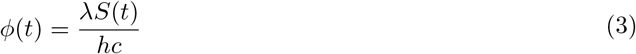

where, S(t) denotes the irradiance of the visual stimulus at time *t, λ* is the wavelength of the visual stimulus, *h* is Planck’s constant, and *c* is the speed of light.

*G*_*d*1_ and *G*_*d*2_ represent the deactivation rates governing transitions from open to closed states, and *G*_*r*_ denotes the thermally driven transition from *C*_2_ to *C*_1_. Because these transitions occur in the absence of light and reflect the return of the opsin to its resting conformation after brief activation, they are treated as constant parameters that are intrinsic biophysical properties of the opsin.

We use the model parameters fitted to photocurrent recordings of ChR2, ReaChR, and ChRmine opsins expressed in rodent retinal ganglion cells as reported in^20^. A summary of these parameters is given in Extended Data (Table 2).

Assuming the visual stimulus is delivered at the opsin’s peak absorption wavelength, the resulting time course of the opsin-mediated photocurrent is expressed as:

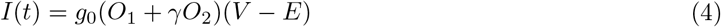

Here, *g*_0_ accounts for both the maximum single-channel conductance and opsin expression density. *V* denotes the membrane potential, held at –60 *mV* during conventional whole-cell patch-clamp measurements, and *E* is the reverse potential of the opsin channel. *γ* specifies the ratio of conductances between the *O*_1_ and *O*_2_ open states.

We simulated opsin-mediated photocurrents *I*(*t*) by numerically solving the ODE system in Eqns. 1-4 using Euler’s method with a *dt* = 0.1 ms timestep. At *t* = 0, we assumed that all the opsin molecules were in closed state *C*_1_. That is, the initial conditions were set to *C*_1_ = 1, *O*_1_ = *O*_2_ = *C*_2_ = 0.

Opsin responses were computed for both movies (shot at 120 fps, upsampled to 840 fps) in Fig. 2, and for sinusoidal visual stimuli of varying spatial and temporal frequencies with a peak irradiance of 1 *mWmm*^−2^ at the chosen opsin’s peak absorption wavelength (460nm for ChR2; 590nm for ReaChR and ChRmine).

In all simulations, to ensure that responses reflected steady-state dynamics rather than adaptation transients, each modeled stimulus was preceded by a 2*s* period of constant illumination at the mean irradiance of the stimulus (0.5 *mWmm*^−2^) for a sinusoid ranging from 0 to 1 *mWmm*^−2^. This pre-stimulus adaptation step approximates retinal adaptation to background illumination.

### 2 Temporal Contrast Sensitivity Measurements

#### Participants

Six human participants (3 male and 3 female), aged 18 to 32 were screened for 20/20 or corrected-to-20/20 vision and had no history of visual impairments. All experiments adhered to ethical guidelines per the approval of the Institutional Review Board of the University of Washington. All participants gave informed written consent prior to the experiment and were informed that they could withdraw at any time.

#### Apparatus

The experiment was conducted in a dark room. The experiment was run on an XPS 13 Plus desktop system with a 12th Gen Intel Core i7-1260P installed with Ubuntu 22.04 LTS OS. All code was written in MATLAB using PsychtoolBox-3 software. Stimuli were presented on a high-resolution (1080pix) on a 32” linearized Display++ monitor with 10-bit depth and a maximum refresh rate of 120 Hz. Participants were seated in a comfortable chair at a distance of 206 cm from the monitor. A chin rest fixed to a resting table was used to e nsure that the participants’ central vision was aligned with the center of the screen. They were provided with a wireless Bluetooth keyboard to record their responses.

#### Stimuli

The stimuli consisted of counterphase sinusoidal Gabor gratings in grayscale of varying spatial frequency (SF) and contrast levels flickering at different temporal frequencies (TF). The SFs and TFs consisted of 12 discrete levels ranging from 0.5 to 16 cpd and 1.5 to 20 Hz respectively. These spatiotemporal sinusoids were windowed with a spatial Gaussian window of *σ*=1 and a temporal ramp function.

We measured tCSFs for four conditions: **(i) Neurotypical/Baseline condition:** unmodified spatially and temporally modulated sinusoidal gratings stimuli, and **(ii)-(iv) ChR2, ReaChR, and ChRmine conditions:** Visual stimuli were filtered using the four-state photocycle model with parameters pertaining to the desired opsin, as described in Section.1, to simulate the temporal dynamics of opsin-mediated vision. We then normalized these absolute photocurrents as shown in (Eqn. 5).

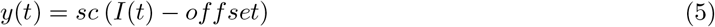

*sc* and *offset* were determined such that the photocurrent response for every opsin produced a steady-state response of 0.5 at the mean luminance of the monitor and peaks at 1 at the lowest temporal frequency in our experiment (1.5 Hz). Scale factors corresponding to the three opsins are reported in Extended Data, Table 3.

Because computing the timecourses was too computationally intensive for real-time rendering, we created a look up table table by pre-computing the photocurrent timecourses for all 1024 possible contrast levels for each of the 12 discrete TFs for each of the three opsins.

#### Experiment

We used a two-alternative forced-choice (2AFC) paradigm to measure tCSFs. Each trial began with a gray screen at the mean luminance of the monitor and a central fixation dot accompanied by a 10 ms, 1000 Hz auditory cue, followed by a blank gray screen. A sinusoidal grating in a cosine ramp window, oriented at either −45^◦^ or +45^◦^, was then presented for 800 ms, followed by a blank gray screen, which remains until a key-press response was made indicating grating orientation via left or right arrow keys. Background contrast is held constant at 0.5 across all trials and conditions (Extended Data, Fig.6).

Grating spatial and temporal frequencies, along with contrast, were selected trial-by-trial using the Bayesian adaptive quick CSF (qCSF) method^44^, which optimizes contrast sensitivity estimates across multiple frequency dimensions through efficient updates to maximize information gain.

The qCSF framework requires fixing spatial frequencies while adaptively presenting stimuli that vary in temporal frequency and contrast, or fixing temporal frequency while adaptively presenting stimuli that vary in spatial frequency and contrast. Therefore, each tCSF measurement consisted of two sessions: (i) fixed SF: five spatial frequencies (0.5, 1.2, 2.8, 6.7, 16 cpd) with adaptively selected temporal frequencies and contrasts, with 50 trials per SF, and (ii) fixed TF: five temporal frequencies (2, 4, 8, 12, 16 Hz) with adaptively selected spatial frequencies and contrasts across 50 trials per TF. Each experiment was repeated five times, yielding 2,500 trials per condition per participant. Sessions lasted about 15 minutes, and condition order was randomized and load-balanced to avoid bias introduced due to order effects, resulting in 10,000 trials across four conditions for each participant. Across six participants, this amounted to a total of 60,000 psychophysical trials.

Baseline contrast detection thresholds corresponding to 20/20 vision (‘neurotypical condition’) were first measured for each neurotypical participant. The experiment was then repeated using the same grating stimuli transformed to simulate optogenetic activation for each of the three opsins (‘ChR2’, ‘ReaChR’, and ‘ChRmine’). The resulting optogenetically driven detection thresholds enabled a direct comparison of perceptual differences in visual acuity attributable to each opsin.

#### Estimation of the tCSF from Psychophysical Data

The qCSF method described above results in trials with stimuli varying non-uniformly in spatial frequency, temporal frequency and contrast. To convert these results into an estimate of the tCSF surface, we partitioned the spatiotemporal region spanning TFs from 0-20 Hz and SFs from 0-16 cpd into a logarithmically spaced 12-by-12 grid. The grid resolution can be adjusted to provide a sufficient number of trials per grid. For each grid cell, we extract the psychophysical trials from each participant and condition (‘Baseline’, ‘ChR2’, ‘ReaChR’, and ‘ChRmine’) whose stimulus frequencies fall within a square window of unit-area in log units from the cell’s center (*sf, tf*). A Weibull function was then fitted to these data using a maximum likelihood cost function to estimate the contrast required to reach a performance level of 80% correct. Because each grid cell contains only about 9 trials, the slope was fixed at 1, leaving threshold as the sole free parameter of the Weibull function.

We repeated this fitting procedure systematically across the entire range of sampled spatiotemporal frequency grids. Contrast sensitivity was defined as the log of the reciprocal of the minimum contrast required to reach 80% correct performance.

One additional step was necessary: because our virtual patients possess a neurotypical retina, their photoreceptors introduce an additional processing stage when viewing opsin-modified stimuli, which does not exist in optogenetic patients. We therefore corrected the estimated tCSFs by compensating for the losses in temporal sensitivity resulting from processing in the outer retinal layers, as described below.

#### Comparing tCSFs across opsins

During human-in-the-loop psychophysics, the grating stimuli for each opsin were scaled such that the difference between the mean and the maximum response for the lowest temporal frequency was the same across all opsins. An average tCSF was then computed across participants for each condition, as shown in Fig. 3D. To isolate the perceptual effects attributable to opsin kinetics, in Fig. 3E we show tCSFs normalized by their relative sensitivity differences at the lowest temporal frequency.

### 3 Predicting clinical outcomes directly from opsin *ex vivo* kinetics

To directly estimate tCSFs from opsin kinetics, we used the opsin photocurrent responses for each temporal frequency to estimate the overall strength of the response by assuming that detection is based on the integration of the response over time, with differential weighting of increments and decrements. This model reduces each corrected opsin-current trace *y*(*t*) to a scalar response

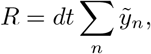

where

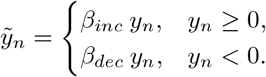

Thus,

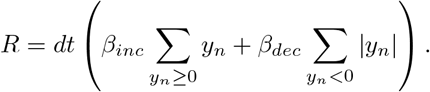

The values of R across temporal frequency, *R*_*k*_, are only relative; thus to predict our psychophysical tCSFs, *R*_*k*_ was scaled according to

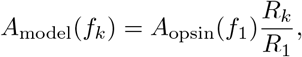

where *A*_opsin_(*f*_1_) is the estimated opsin attenuation at the lowest temporal frequency, and *R*_1_ is the response of the model at the lowest temporal frequency.

Finally, the opsin-modified tCSF was predicted by:

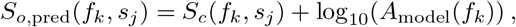

where *s*_*j*_ indexes spatial frequency.

The procedure involves integrating the time-course of the opsin response while weighting positive and negative components differently. Estimates of the positive components, *β*_*inc*_, and the negative components, *β*_*dec*_, for each spatial frequency were obtained by minimizing the mean squared error between the psychophysical measurements corresponding to the three microbial opsins and model predictions of sensitivity. Importantly, the same values of *β*_*inc*_ and *β*_*dec*_ fit all three opsins, allowing this model to be generalized to novel opsins without the need for additional data (see Extended Data, Table 4).

### 4 Compensating for Outer Plexiform Processing

The temporal sensitivity of our human-in-the-loop virtual patients is determined by two factors: (i) the simulated latency in opsin-generated photocurrents, and (ii) the additional processing resulting from the fact that our virtual patients are viewing these modified stimuli through their own, intact, outer plexiform layer. Thus, tCSF estimates derived from human-in-the-loop virtual patients must be corrected to account for intrinsic dynamics of the outer plexiform layer.

Under steady (constant) mean luminance and for small-signal modulations within our limited temporal-frequency range, the temporal dynamics of the outer plexiform layer can be approximated as a linear, time-invariant filter^45^.

Assuming linearity of the outer plexiform layer of the retina, the retina response *r*(*t*), can be modeled as:

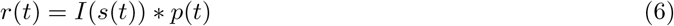

where, *s*(*t*) represents the original sinusoidally modulated stimulus over time, *I*(*t*) is the photocycle model for a given opsin, and *p*(*t*) is the impulse-response function of the outer plexiform layer. The ∗ operator denotes convolution.

We estimated the human cone photocurrent impulse response *p*(*t*) using the van Hateren phototransduction model^27^ (assuming a 20mm pupil). By converting *p*(*t*) into the Fourier domain, *P* (*f*), the effect of the outer plexiform layer can be corrected for by simply multiplying the amplitude of *I*(*s*(*t*)) by *P* (*f*).

Figure 4 compares the original tCSFs (solid colors, same as in Fig. 3E) to the tCSFs after correcting for the effects of the outer plexiform layer (Fig.4, translucent colors). The frequency response cuts off drastically around 54 Hz for neurotypical human retina. Therefore, we do not observe a significant difference between the original and adjusted tCSFs in the range of frequencies we tested; the effects of the dynamics of the outer plexiform layer on the TCSF were negligible compared to the delays due to opsin kinetics. However, photoreceptor correction may prove to be more important in the future for faster opsins.

## Extended Data

**Fig. 6:**
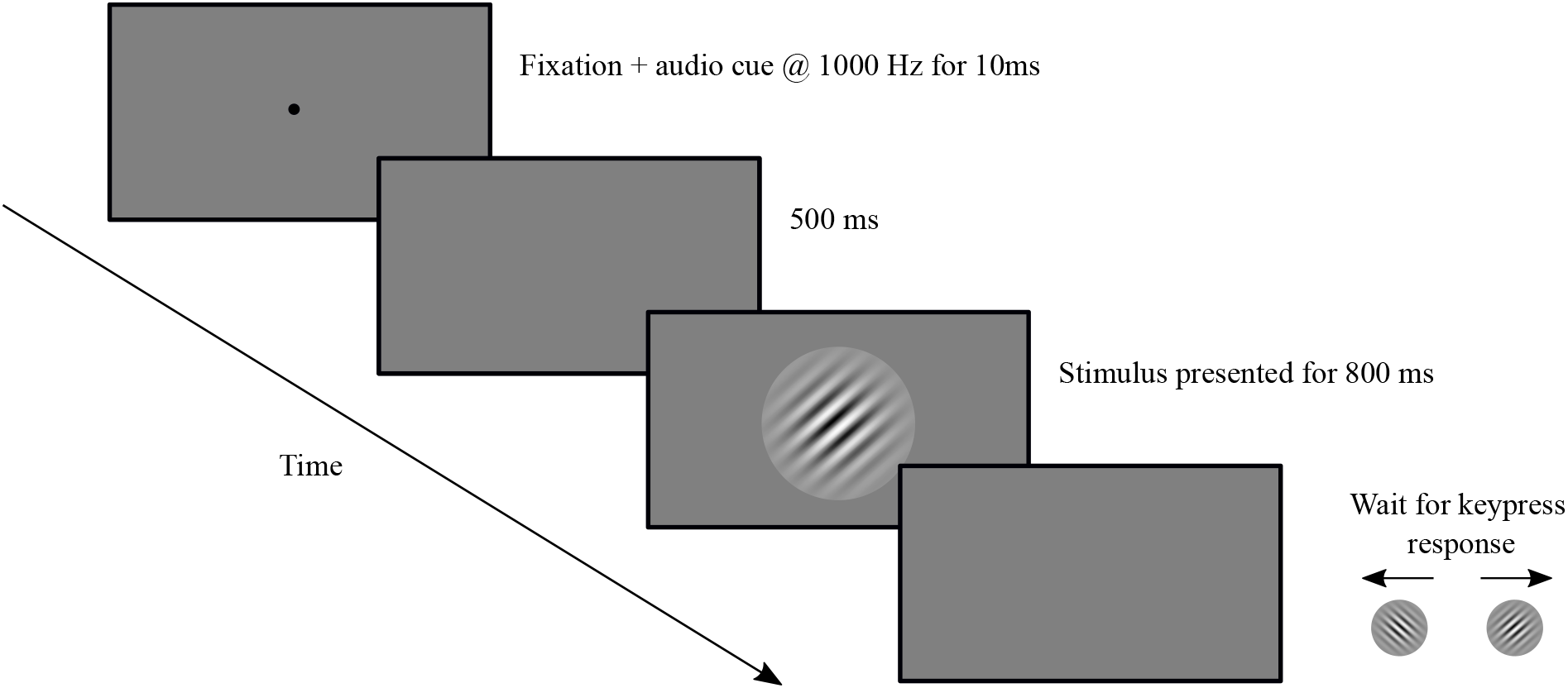
Measuring contrast sensitivity function with a 2AFC orientation discrimination task. An audio cue of 1000 Hz, accompanied by a central fixation dot, signals the trial onset to the virtual patient. 10 ms later the stimulus appears for 800 ms, oriented either at +*/* −45^*o*^ degrees and varying in spatial and temporal frequency across trials. The display then switches to a uniform gray screen, which remains until a keypress response is made. Background contrast is held constant at 0.5 across all trials and conditions.

**Fig. 7:**
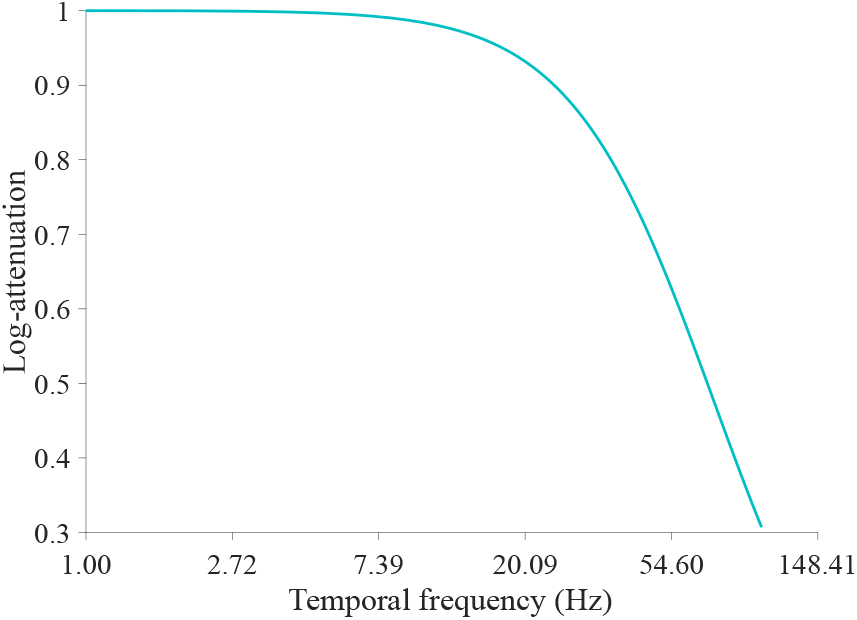
Frequency response of human cone photoreceptors. Photocurrent responses were modeled using the Van Hateren phototransduction model^27^ for sinusoidal gratings of unit amplitude and varying frequency. Log-attenuation was computed as the log of the ratio of peak amplitude at a given frequency to that of the lowest temporal frequency.

**Table 3.** Scale factors required to generate normalized optogenetic timecourses.

| Opsin | sc | offset |
| --- | --- | --- |
| ChR2 | -41.89 | 0.0258 |
| ReaChR | -235.19 | 0.0064 |
| ChRmine | -1147.80 | 0.0012 |

**Table 4.** Parameter estimates for our linking hypothesis model.

| SF (cpd) | $\beta_{\text{inc}}$ | $\beta_{\text{dec}}$ |
| --- | --- | --- |
| 0.3 | 0.6066 | 1.1951 |
| 8.03 | 0.8648 | 0.7590 |
| 16.72 | 0.9021 | 0.6898 |

## Declarations

## Acknowledgments

We sincerely thank Vijayakrishna Naganoor and Vasishta Divakar Harekal for their assistance in creating the simulation videos. We also thank Dr. Kelly Chang for helpful discussions regarding the manuscript. This work was supported by the National Eye Institute (grant number EY031312 to I.F. and G.M.B.). This work was facilitated through the use of advanced computational, storage, and networking infrastructure provided by the Hyak supercomputer system at the University of Washington.

## Code Availability

All code pertaining to the virtual patient software, generating simulations, and data analysis used in this study is publicly available on GitHub at https://github.com/Vaishnavi-B-Mohan/p2poptoExpt.

## Data Availability

The raw human psychophysics data generated in this study are not publicly available due to institutional restrictions regarding participant privacy and ethical consent. However, de-identified data are available from the corresponding authors upon request and subject to approval by the University of Washington Institutional Review Board. The full supplementary videos of the simulations are available from the corresponding authors upon request.

## Author contribution

All authors contributed to conceptualization and methodology. V.B.M., I.F., and G.M.B. contributed to writing. V.B.M. and G.M.B. contributed to software.

## Ethics approval and consent to participate

All experimental procedures involving human participants were approved by the Institutional Review Board at the University of Washington. All participants provided written informed consent prior to engaging in the psychophysics experiments.

